# Diet and injection, our recommendation to characterize *Clavibacter michiganensis*- tomato interactions

**DOI:** 10.1101/2023.04.28.538756

**Authors:** Anne-Sophie Brochu, Jeanne Durrivage, Dagoberto Torres, Edel Pérez-López

## Abstract

Tomato (*Solanum lycopersicum* L.) is one of the most important vegetables in the world. Its extensive cultivation has made this plant the target of many viral, fungal, and bacterial diseases. Among them, the bacterial canker of tomato caused by *Clavibacter michiganensis* (*Cm*) has been named one of the most devastating diseases affecting the tomato industry worldwide. It can significantly reduce the yields and profitability of this crop. One of the big challenges we found when working with *Cm* and trying to characterize the virulence of different isolates was the lack of a consensus methodology to inoculate tomato plants, how to fertilize them and characterize *Cm* virulence. This research aimed to identify an artificial inoculation method to induce bacterial canker on tomato plants in greenhouse conditions to homogenize the results of different studies with *Cm*. We compared two inoculation methods, including the scalpel and syringe method with two levels of fertilization, low and high fertilization. After evaluating several variables like the percentage of necrotic leaves and the height of the plants, the results showed that the syringe inoculation with low fertilization was the most effective inoculation method allowing the development of a multilevel scale that can be used to study the interaction between tomato plants and *Cm* isolates.

## INTRODUCTION

Tomato (*Solanum lycopersicum* L.) is one of the most consumed vegetables in the world with a global production of more than 189 million tons of over 5 million hectares (FAO, 2021). The United States was the world’s second-largest producer of tomatoes in 2021 with a production of over 11 million tons and revenues of over $1.5 billion (FAO, 2021; USDA, 2022). However, tomato is susceptible to many diseases such as bacterial canker of tomato, caused by the gram-positive bacterium *Clavibacter michiganensis* (*Cm* hereafter) belonging to the class actinomycetes (Davis et al., 1984; Li et al., 2018). It is one of the most devastating bacterial diseases affecting the tomato industry worldwide causing considerable yield losses (Chang et al., 1992; Emmatty and John, 1973; Poysa, 1993; Nandi et al., 2018). *Cm* was first reported in the 1900s in Michigan and since then it has caused several outbreaks in most tomato-growing regions (Kleitman et al., 2008; Blank et al., 2016; Boyaci et al., 2021). *Cm* mainly affects, tomatoes, and causes marginal necrosis, canker formation, plant wilting and brown discolouration of vascular tissues (Gleason et al., 1993; Chalupowicz et al., 2012; Sen et al., 2015). This pathogen invades vascular tissues usually xylem through wounds or natural openings such as stomata and hydathodes (Carlton et al., 1998; Tancos et al., 2013) which allows the disease to be spread by splashes or drops of water or contaminated tools (Chang et al., 1991; Gitaitis et al., 1991; Carlton et al., 1998; Xu et al., 2010).

In Quebec, Canada, the greenhouse industry is experiencing an unprecedented expansion, and with it, is expected an increase in phytosanitary problems (Wheeler 2023). After the tomato brown rugose fruit virus, the bacterial canker of tomato is considered the second-biggest threat to greenhouse production worldwide, reason why growers and the industry are actively looking for resistant germplasm or solutions compatible with organic production (Abo-Elyousr et al., 2019; Abebe et al., 2022). A challenge presented to us by several industry partners was how to obtain a homogenous infection with *Cm* to be able to study the response of different tomato varieties to the disease or to test the efficiency of new environmental-friendly biological control agents or disinfection methods.

Different inoculation methods of *Cm* have been used for tomato cultivar resistance evaluation, to study the virulence of certain isolates, and to study the efficiency of control methods against the pathogen (Bulk et al., 1991; Kleitman et al. 2008; Takishita et al. 2018). In most cases, the concentration of the inoculum, the phenological stage of the tomato plant during the inoculation, growth conditions, how the symptoms are evaluated, and the time of evaluation differed from one study to another impossible to compare them. In this study, we considered the protocols employed by various research groups and compared two of the most popular inoculation methods to identify the most efficient. In addition, we investigated how the level of fertilization could contribute to better characterizing the physiological response of tomato plants to *Cm* infection.

## METHODOLOGY

### Bacterial isolate and inoculum preparation

Pathogenic isolate *Cmm-1375*, isolated from Quebec, Canada, and provided by the provincial plant pathology diagnostic laboratory (LEDP, from French *Laboratoire d’expertise et de diagnostic en phytoprotection*) was used in this study. Liquid cultures were prepared using BD Bacto™ Tryptic Soy Broth (Soybean-Casein Digest Medium). Bacterial cultures were prepared from glycerol stock stored at -80°C with a final concentration of 20% glycerol with an inoculation loop of 10 μl in Erlenmeyer flask of 50 ml of TSB medium. The inoculated TSB media were incubated at 28°C and 200 rpm for 2 days. Bacteria were subsequently pelleted by centrifugation at 3500 g for 5 min and washed in 10 sterile mM MgCl_2_ twice. The bacterial solution was diluted with 10 mM MgCl_2_ to an OD_600_ of 0.1, approximately 10^8^ CFU/ml (Epoch 2 Microplate Spectrophotometer) (Poysa, 1993; Kleitman et al., 2008; Thapa et al., 2020; Chalupowicz et al., 2017) and was used immediately to inoculate tomato plants.

### Plant growth

The tomato cultivar M82, previously identified as susceptible to *Cm* (Thole et al., 2021) was grown under greenhouse conditions at 26°C during the day and 18°C at night with 60% relative humidity and a 16 h photoperiod. Seeds were planted in 209 Tray Pallet (diameter: 20 mm; length: 30 mm) filled with PRO-MIX BX (Premier Tech Horticulture, CAD) previously moistened. A plastic dome was added to cover the seedlings until germination to conserve moisture. Two weeks after planting, seedlings were transplanted into 5″ pots filled with PRO-MIX BX (Premier Tech Horticulture, CAD). Plants were watered with the nutrient solution once, containing 0.22 g/L of 15-0-0 +19% calcium fertilizer (Plant-Prod, CAD) and 0.29 g/L of 6-11-31 fertilizer (Plant-Prod, CAD) until the water saturation point of the substrate was reached. The pH was adjusted to 5.5–6.0 with phosphoric acid 75% solution (Plant-Prod, CAD). All the plants were watered as needed until 1 day before inoculation. Three-week-old tomato plants, at the 3-leaf stage, were used for all inoculations.

### Inoculation methods

To identify the most effective inoculation method, two methods were evaluated, syringe and scalpel. For syringe inoculation, three-week-old plants were wounded by removing a cotyledon and the injection was performed using a 1 ml syringe HENKE-JECT (VWR, CAD) and a needle 26G 12.7 mm (1/2”) (VWR, CAD) with 10 μl of bacterial solution with an OD_600_ of 0.1 (Epoch 2 Microplate Spectrophotometer). For scalpel inoculation, three-week-old plants were also wounded by removing a cotyledon and cut with a scalpel previously soaked in the bacterial solution with an OD_600_ of 0.1 (Epoch 2 Microplate Spectrophotometer) (Fig. S1). Tomato plants used as a control in both methods were inoculated with 10 mM MgCl_2_ following a similar procedure.

### Fertilization

After the inoculation, plants were fertilized using two different concentrations. High fertilization treatment received fertilization in two out of three irrigation, and low fertilization was received in one out of three irrigations (Fig. S1). The fertilization was composed of 0.44 g/L of 15-0-0 +19% calcium fertilizer (Plant-Prod, CAD) and 0.58 g/L of 6-11-31 fertilizer (Plant-Prod, CAD). The pH was adjusted to 5.5–6.0 with phosphoric acid 75% solution (Plant-Prod, CAD). After the inoculation, plants were kept in the greenhouse in the same condition as mentioned above. The plants were watered at the same time with or without fertilization depending on fertilization treatment as needed and until the water saturation point of the substrate was reached. The experiment was set up according to a generalized randomized block design with two repetitions in time of six plants per treatment for each of the eight combinations of inoculation methods, fertilization levels, and inoculum.

### Foliar disease symptoms and growth evaluation

Disease evaluation was based on bacterial canker symptoms. The number of leaves of the plants and the number of leaves with necrosis were counted at 7, 14 and 21 days after inoculation (dpi) to obtain the percentage of necrotic leaves per plant. Based on the percentage of necrotic leaves, a disease severity scale for assessing leaf disease symptoms in this study was developed. Plant height was also measured at 7, 14 and 21 dpi to assess the influence of fertilization and the impact of the pathogen on plant growth.

### DNA extraction and multiplex PCR assay

Plant tissue was harvested at 21 dpi from 3 plants of different treatments. To obtain DNA from inoculated plants and control plants, 1 cm stem fragments were collected at the inoculation site, 3 and 6 cm above the inoculation site and from the rachis of the third leaf. The samples were dried using a freeze dryer and macerated in a 2 ml tube. Then, grounded tissue was used to extract genomic DNA using the cetyltrimethylammonium bromide (CTAB) method as previously described (Murray and Thompson 1980). To confirm the presence of *Cm* in each tissue, we use the multiplex PCR assay to detect pathogenic *Cm* (Thapa et al., 2020). The amplification products were observed by electrophoresis using 1 % agarose gel stained with SYBR Safe (ThermoFisher, CAD) and visualized under UV light.

### Statistical analysis

The effects of inoculation methods, fertilization levels, and time on the percentage of necrotic leaves were studied using a binomial generalized linear mixed model. In this model, the empirical logit function was used as the link function, instead of the traditional logit function, because the latter is not defined for percentage values of 0% or 100%. Based on the AIC criterion, the compound symmetry correlation structure was chosen to account for the dependence between observations taken over time on the same experimental unit. Negative controls for each inoculation method and fertilization level did not show symptoms and were not included in the model. In addition, the percentage of necrotic leaves at 7 days was removed from the data set due to the low presence of symptoms. In this model, inoculation methods (2 levels), fertilization levels (2 levels), and time (2 levels), as well as all two- and three-way interactions were considered as fixed factors, whereas plants and replicates were used as random factors.

In the plant height analysis, the inoculum factor and interaction terms involving this factor were added to the model as fixed terms, since negative controls were considered this time. However, the analysis was performed by fitting a linear mixed model on the data since the response variable is continuous. With this model, the AR(1) correlation structure now fits the data best. The analysis was performed by transforming the height measures by the square root, as suggested by the Box-Cox technique, to satisfy the normality assumption.

In both analyses, following a significant effect of one source of variation, we used Tukey’s multiple comparisons method to identify the levels of that source that differed from the others, while controlling for the type I error rate. All analyses were performed using R software version 4.2.3 (R Core Team, 2023) with the lme4 package glmer function for the binomial generalized linear mixed model, and the nlme package lme function for the linear mixed model, at the 0.05 significance level. The code and raw data used for the analysis can be found here: https://github.com/Edelab/Clavibacter-michiganensis-inoculation-method.

## RESULTS

### Scalpel vs syringe

#### Necrotic leaves

Mock-inoculated plants for each inoculation method and fertilization level were symptomless and were not included in the percentage of necrotic leaf analyses. The percentage of necrotic leaves at 7 days was also removed from the data set due to the low presence of symptoms. The percentage of necrotic leaves was significantly different between the ‘scalpel’ and the ‘syringe’ inoculation methods (*p* < 0.001). Using syringe inoculation, more necrotic leaves were observed than with scalpel inoculation at 14 dpi and 21 dpi (Fig. 1A, Table S1). There was also a significant two-way interaction between fertilization level and timing observation (*p* = 0.0140). More necrotic leaves were observed at 21 dpi than at 14 dpi, with no significant difference between the low and high fertilization treatments (Fig. 1B-C). However, the increase in the percentage of necrotic leaves between 14 dpi and 21 dpi is more pronounced in plants with low fertilization treatment (Fig. 1D)

**Figure 1:**
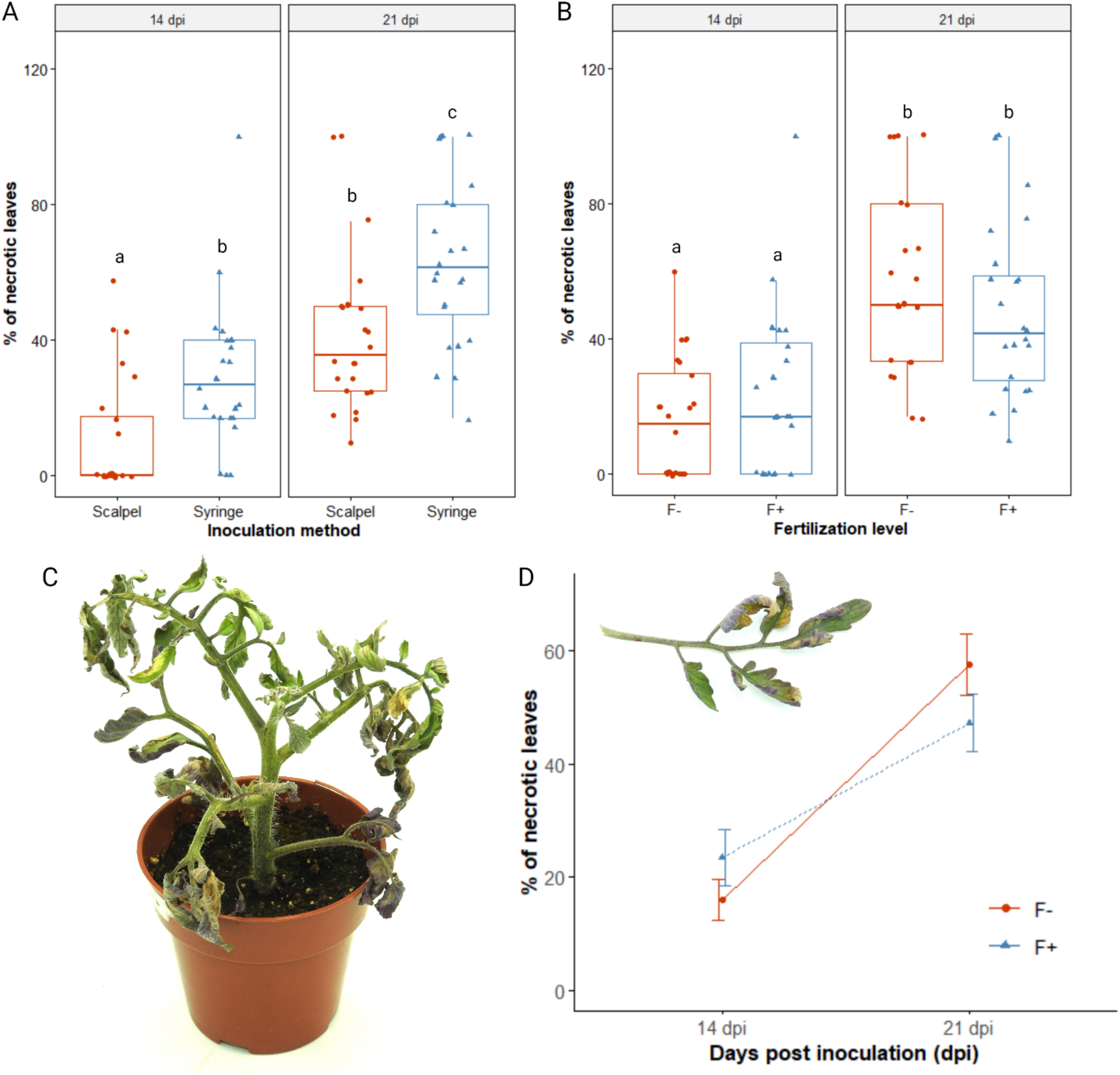
Percentage of necrotic leaves. A, in two different inoculation methods at 14 and 21 dpi. B, in two different fertilization levels at 14 and 21 dpi. Red = Scalpel; Blue = Syringe; Different letter = statistically significant difference (*p* < 0.05) in the proportion of necrotic leaves, according to the generalized linear mixed model, n=12 per treatment. C, tomato plant affected by *Clavibacter michiganensis* at 21 dpi. D, comparison of the percentage of necrotic leaves at 14 and 21 dpi under low (red) and high (blue) fertilization regimes.

#### Plant height

When analyzing how the infection was influencing plant height, a three-way interaction between inoculum, inoculation methods and observation time is significant (*p* = 0.0098) (Table S2). At 7 dpi, all inoculation methods, mock and *Cm*-inoculated plants had similar heights (Fig. 2). At 14 dpi, mock-inoculated plants have similar heights, whereas *Cm*-inoculated plants under both inoculation methods have a lower height than mock plants. Plants inoculated using a syringe presented a significantly lower height than those inoculated with a scalpel, with differences observed at 14 and 21 dpi (Fig. 2A, C). This difference was also observed between infected plants and mock-inoculated control (Fig. 2A, C). A three-way interaction between inoculum, fertilization levels, and observation time was also significant (*p* = 0.0098). At 7, 14, and 21 dpi, a significant difference in height is observed between the two fertilization treatments (Fig. 2B, C). Generally, we observed that plants with higher fertilization were taller, either mock- or *Cm*-inoculated (Fig. 2B, C, Table S2). However, the low fertilization treatment is not significantly different between 14 dpi and 21 dpi.

**Figure 2:**
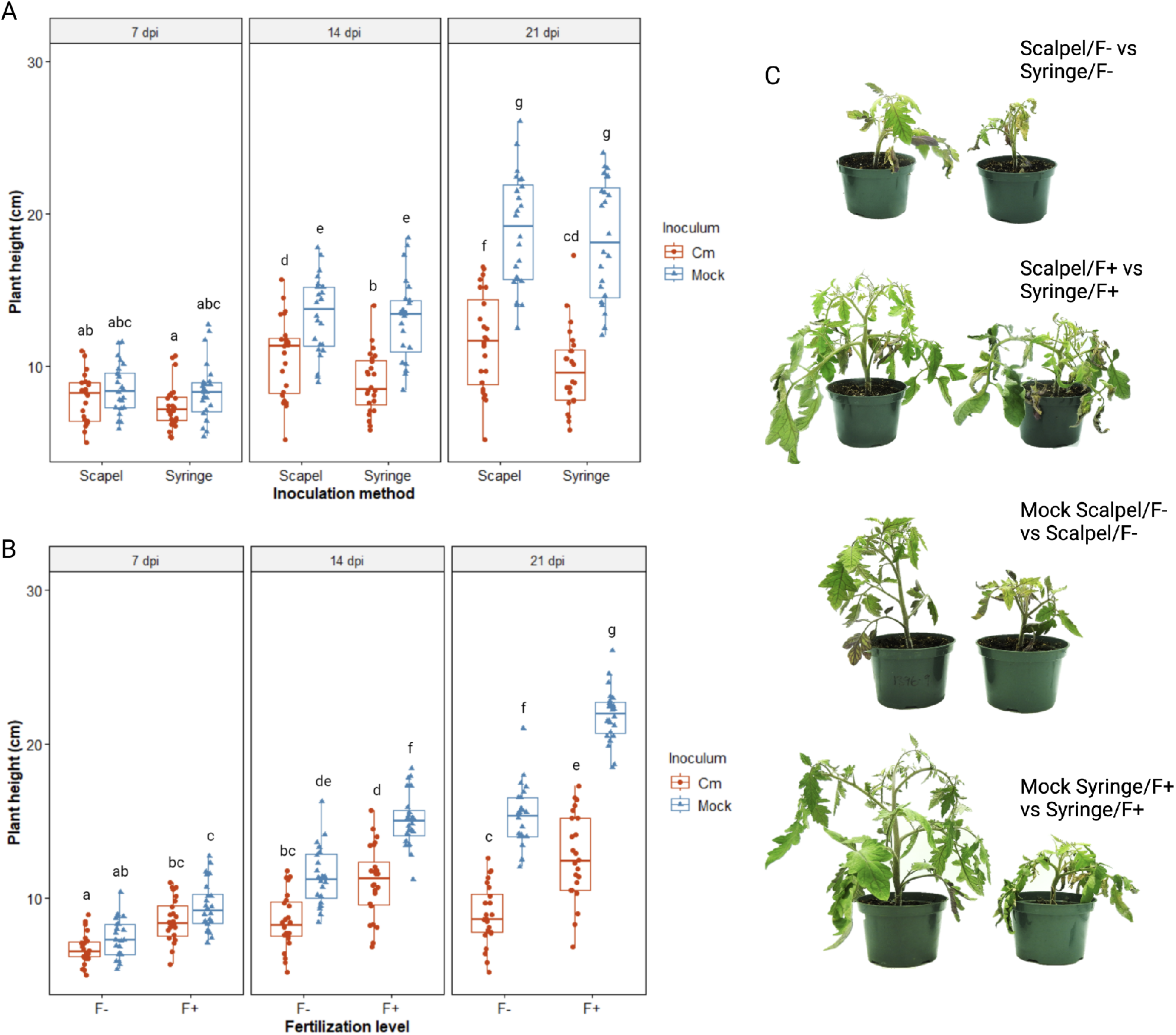
Plant height. A, of *Clavibacter michiganensis* (*Cm)*- and mock-inoculated plants using scalpel or syringe inoculation at 7, 14 and 21 dpi. B, of *Cm*- and mock-inoculated plants using low and high fertilization levels at 7, 14 and 21 dpi. Red = *Cm-*infected plants; Blue = Mock plants. Different letters = statistically significant difference (*p* < 0.05) in the proportion of necrotic leaves, according to the generalized linear mixed model, n=24 per treatment. C, representation of tomato plant phenotype under the different treatments investigated in this study.

For every treatment *Cm* was detected in all *Cm-*infected plants using the multiplex PCR assay previously developed by Thapa et al. (2020) (Fig. S2). The three amplicons belonging to *rhuM* (1 000 bp), *tomA* (630 bp), and *16S rRNA* (415 bp) were amplified from *Cm-*infected plants, while only the fragment corresponding to 16S rRNA was amplified from healthy plants (Fig. S2), confirming that independently of the inoculation method and fertilization regime, *Cm* can reach the same parts of the plant.

### A score scale to characterize *C. michiganensis-*tomato interactions

The first symptoms of *Cm* infection observed on the M82 tomato plants became visible at 7 dpi for the two inoculation methods and fertilization levels. The main symptoms observed were yellowing of the older leaves followed by wilting, necrosis on the leaves, and stem canker. After evaluating all the plants under the different treatments, we concluded that the best method to inoculate *Cm* was using a syringe and keeping the inoculated plants under a low fertilization regime. Those were the conditions used by us to design a rating scale to calculate the bacterial canker of tomato disease index (Fig. 3) (Chiang et al., 2017). For that, we propose the use of the percentage of necrotic leaves as a quantitative variable and a scale from 0 to 5, where 0 represents no infection and 0% of necrotic leaves and 5, the highest level of infection and 75-100% of necrotic leaves (Fig. 3).

**Figure 3:**
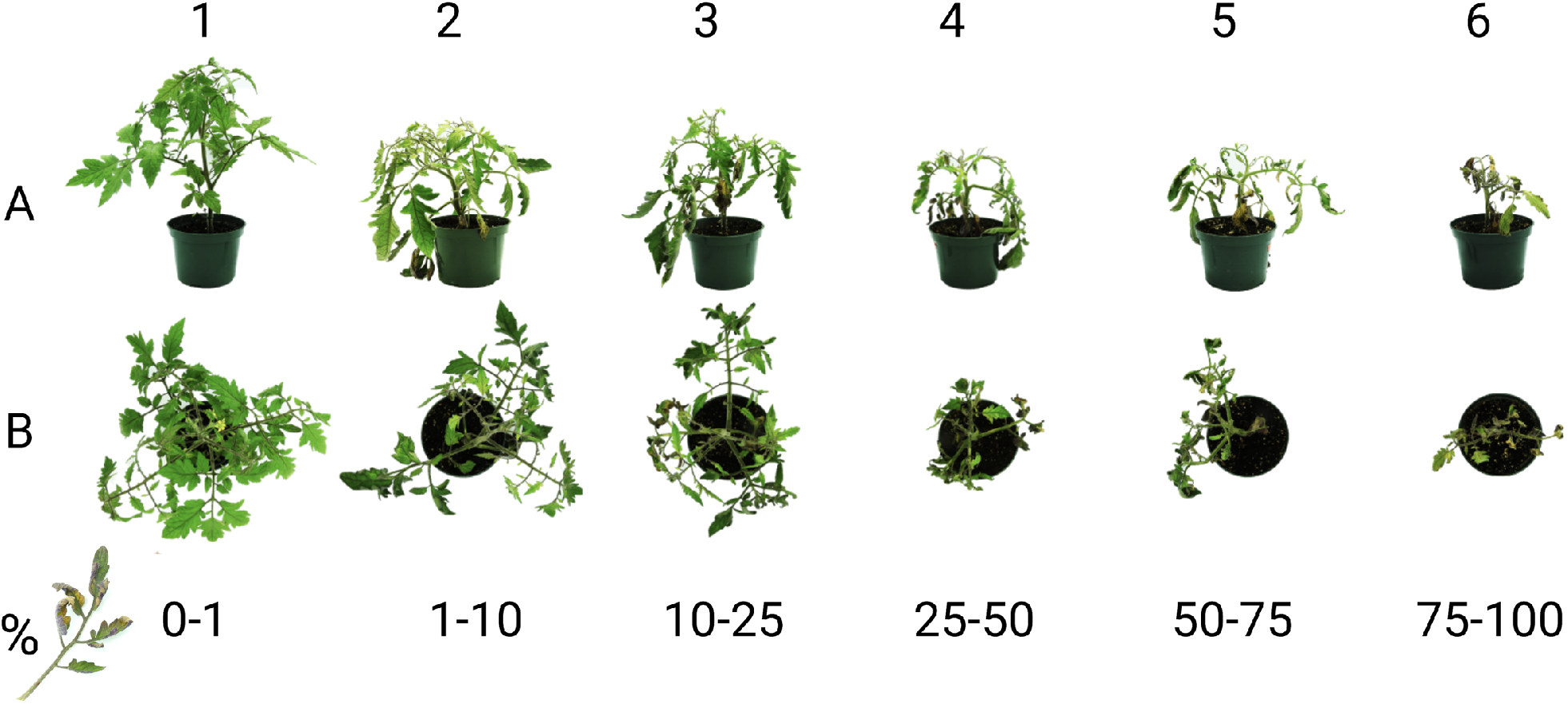
Symptoms score scale to evaluate tomato plant response to *Clavibacter michiganensis*. A, tomato plants are seen from the side. B, tomato plants are seen from above. The percentage of necrotic leaves for each level is presented below. For the preparation of this scale, we used the bacteria isolated *Cmm-1375* and the tomato cultivar M82.

## DISCUSSION

Several inoculation methods have been used by different research groups. The main methods used are the scalpel or scissors previously dipped in bacterial suspension (Bulk et al., 1991; Poysa 1993; Francis et al., 2001; Kaneishiro et al.,2006; Mohd Nadzir et al., 2019), the root-dipping in bacterial suspension (Poysa 1993; Abo-Elyousr et al., 2019), stem injection of the bacterial suspension with a syringe (Milijaševic-Marcic et al., 2012; EPPO, 2016; Osdaghi et al., 2018; Takishita et al., 2018), the soil/media drench method (Mohd Nadzir et al., 2019; Oloyede et al., 2021), and the application of a bacterial solution to a wound (Chalupowicz et al. 2017; Thapa et al. 2020). Some studies have compared more than one method, finding, for example, that plants showed more symptoms when inoculated with scissors dipped in bacterial suspension than the roots-dipping method (Poysa, 1993). Similar conclusions were obtained when adding the bacterial suspension to the plant medium compared with direct inoculation by wounding the stem (Mohd Nadzir et al. 2019). This high diversity of methods makes it difficult to compare studies among them and calls for a more homogenous methodology like the one presented by us.

Fertilization can have an impact on the growth and development of tomato plants, which in turn can affect their susceptibility to disease (Nicot et al., 2012). Some studies have shown a decrease in disease severity due to low fertilization, although opposite effects have been also reported (Snoeijers et al., 2000; Fagard et al., 2014). Here we found a decrease in the severity of the disease in those plants under high fertilization by decreasing the proportion of necrotic plants and increasing the height of infected plants, while infected plants under lower fertilization showed more pronounced symptomatology. Plants lacking nitrogen are weaker, grow slower and age faster (Snoeijers et al., 2000). Some studies have indicated that nitrogen fertilization could promote plant colonization by biotrophic pathogens like *Cm* (Snoeijers et al., 2000; Fagard et al., 2014). However, a study focused on *Xanthomonas campestris* pv. *campestris* - cabbage interaction reported that high levels of nitrogen significantly reduced the level of systemic xylem colonization by the bacteria as well as the development of black rot lesions (McElhaney et al., 1998). No study has been conducted on the impact of fertilization on the severity of bacterial canker of tomatoes. However, one study was conducted with the species *Clavibacter spedonicus* showing that higher fertilization increases the severity of the disease (Walker and Gallegly, 1951). Overall, the impact of fertilization on host response to *Cm* is complex and depends on a variety of factors, including plant growth stages, cultivars, type and amount of fertilizer applied, the timing of the application, and environmental conditions (Richard-Molarde et al.1999; Berry et al. 2010).

Historically, wilting has been considered the main symptom when studying bacterial canker of tomato severity, and at least one symptom score scale has been developed to calculate the disease index (Mohd Nadzir et al., 2019; Takishita et al. 2018; Abo-Elyousr et al. 2021). Nonetheless, the determination of the percentage of wilting is subject to the evaluator’s interpretation and has a high level of human error associated with it. That is different for the percentage of necrotic leaves (Fig. 3), a variable easy to quantify and easily reproducible under the conditions suggested in this study.

In conclusion, here we identified the best inoculation and fertilization method to study tomato physiological response to bacterial canker on tomato in greenhouse conditions. In doing so, we were able to generate a new symptom score scale that will facilitate the characterization of tomato germplasm to *Cm* and/or the pathogenicity of *Cm* isolates. The methodology and score scale described here will have great implications for the tomato industry and contribute to keeping the search for a *Cm*-resistant cultivar.

## ACKNOWLEDGEMENTS

The authors would like to thank the Laboratoire d’expertise et de diagnostic en phytoprotection of MAPAQ for providing *C. michiganensis* isolate used in this study. We also thank NSERC for funding this research through the Discovery program granted to EPL and the USRA and Master’s scholarships granted to ASB. We also thank MITACS for supporting Dagoberto Torres through the Globalink program.

---

**Table S1:** Statistic test for the effect of two different inoculation methods and two fertilization levels on the percentage of necrotic leaves of tomato M82 at 14- and 21-days post inoculation

| Effect | Anova table |  |  |
| --- | --- | --- | --- |
|  | df<sub>N</sub> | Chi-square | P value |
| Inoculation method | 1 | 17.439 | <.0001 |
| Fertilization level | 1 | 0.004 | 0.9498 |
| Inoculation method x Fertilization level | 1 | 0.738 | 0.3903 |
| Time | 1 | 66.756 | <.0001 |
| Inoculation method x Time | 1 | 1.715 | 0.1904 |
| Fertilization level x Time | 1 | 6.042 | 0.0140 |
| Inoculation method x Fertilization level x Time | 1 | 0.658 | 0.4174 |

df<sub>N</sub> = numerator degrees of freedom

**Table S2:** Statistic test for the effect of two different inoculation methods and two fertilization levels on tomato M82 plant height at 7-, 14- and 21-days post inoculation.

| Effect | Analysis of variance-type statistics |  |  |  |
| --- | --- | --- | --- | --- |
|  | df<sub>N</sub> | df<sub>D</sub> | ATS | P value |
| Inoculum | 1 | 87 | 191.902 | <.0001 |
| Inoculation method | 1 | 87 | 9.874 | 0.0023 |
| Fertilization level | 1 | 87 | 139.676 | <.0001 |
| Inoculum x Inoculation method | 1 | 87 | 4.383 | 0.0392 |
| Inoculum x Fertilization level | 1 | 87 | 2.387 | 0.1260 |
| Inoculation method x Fertilization level | 1 | 87 | 1.156 | 0.2854 |
| Inoculum x Inoculation method x Fertilization levels | 1 | 87 | 2.399 | 0.1250 |
| Time | 2 | 175 | 1323.212 | <.0001 |
| Inoculum x Time | 2 | 175 | 320.716 | <.0001 |
| Inoculation method x Time | 2 | 175 | 8.271 | 0.0004 |
| Fertilization level x Time | 2 | 175 | 34.936 | <.0001 |
| Inoculum x Inoculation method x Time | 2 | 175 | 4.745 | 0.0098 |
| Inoculum x Fertilization levels x Time | 2 | 175 | 4.492 | 0.0125 |
| Inoculation method x Fertilization levels x Time | 2 | 175 | 2.364 | 0.0970 |
| Inoculum x Inoculation method x Fertilization level x Time | 2 | 175 | 1.995 | 0.1391 |

df<sub>N</sub> = numerator degrees of freedom and df<sub>D</sub> = denominator degrees of freedom.

**Fig. S1:**
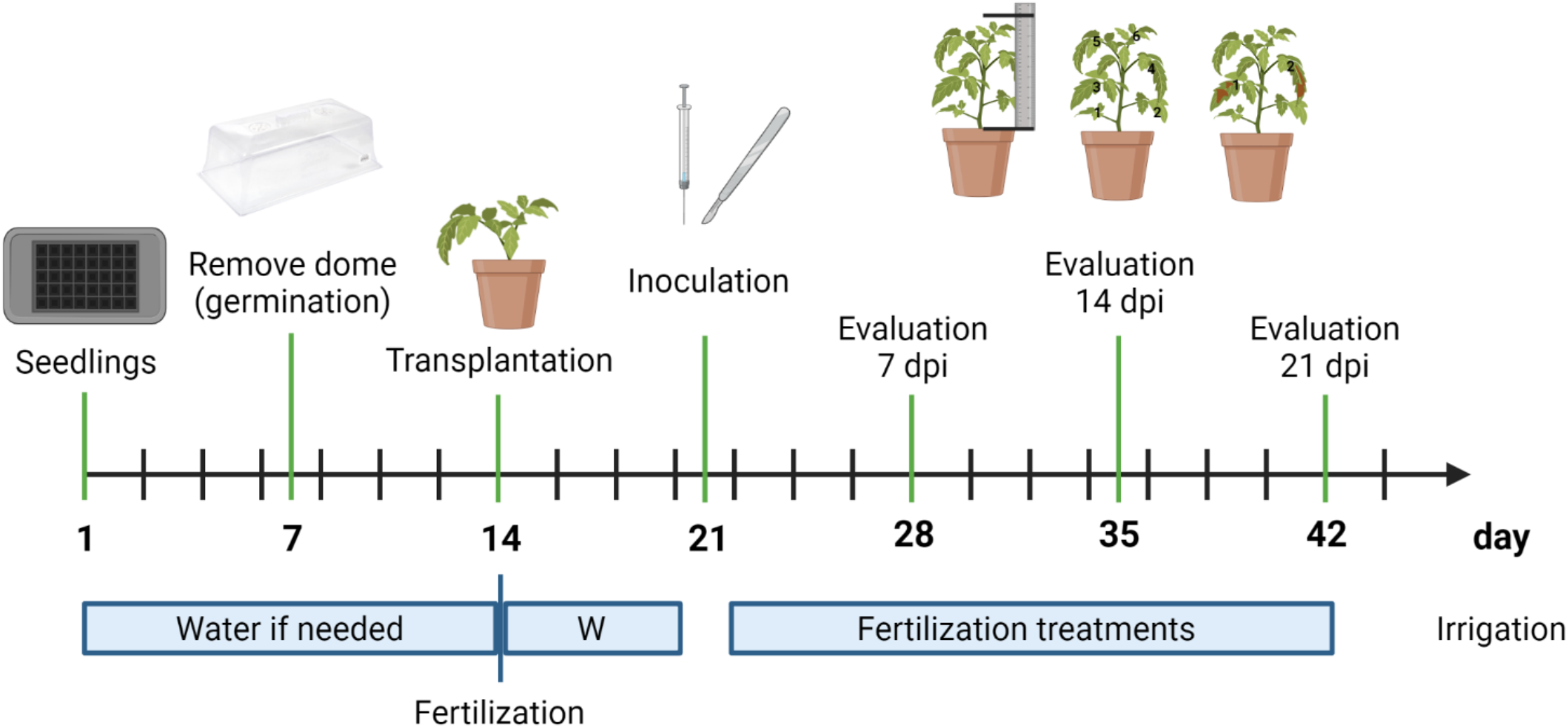
Schematic representation of the methodology used in this study.

**Figure S2:**
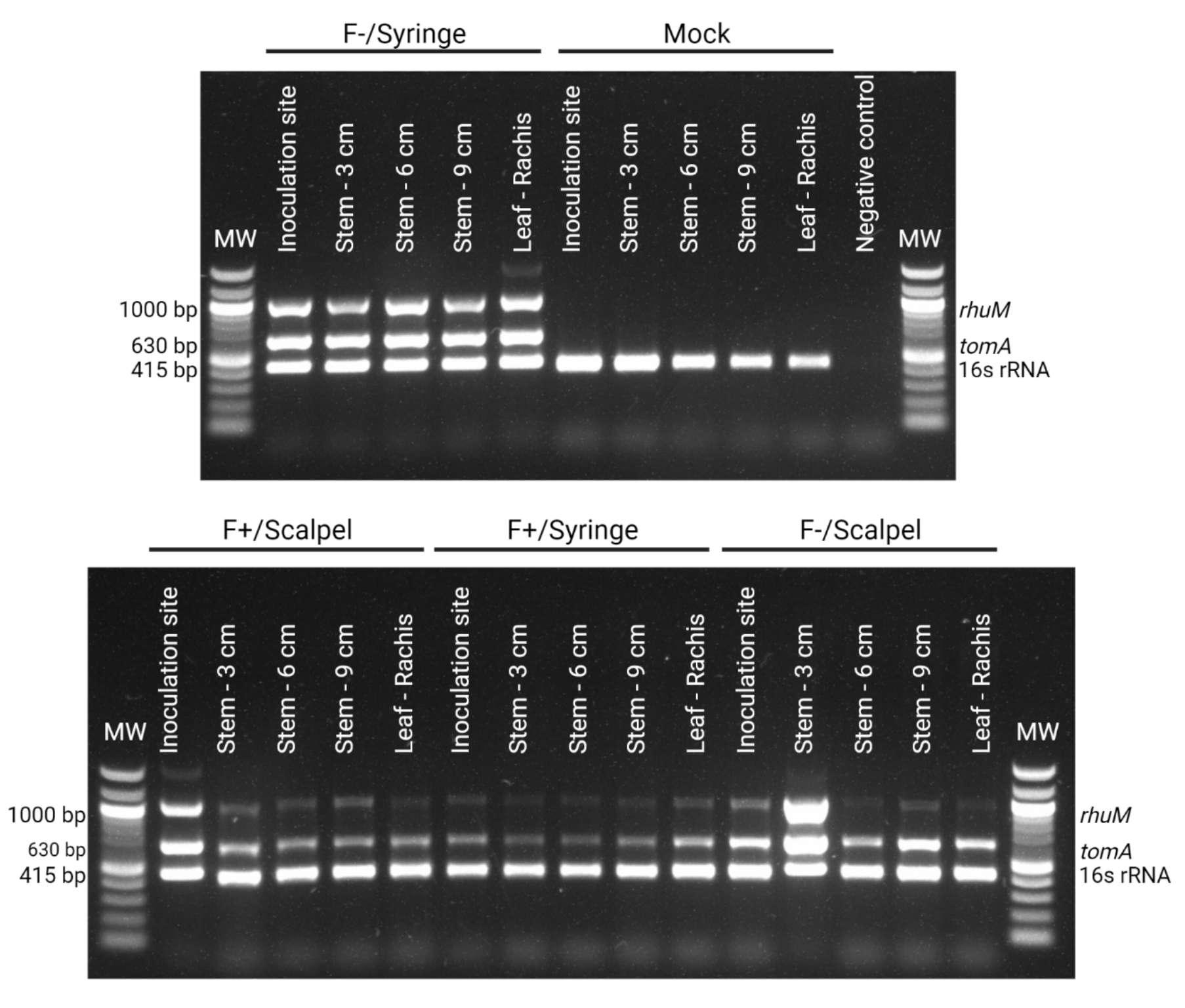
Detection of *Clavibacter michiganensis* using the multiplex PCR developed by Thapa et al. (2020). Agarose gel electrophoresis, MW, molecular weight marker 100 bp DNA ladder.

